## Supplementary_Info for "Increasingly efficient chromatin binding of cohesin and CTCF supports chromatin architecture formation during zebrafish embryogenesis"

Supplementary Figures

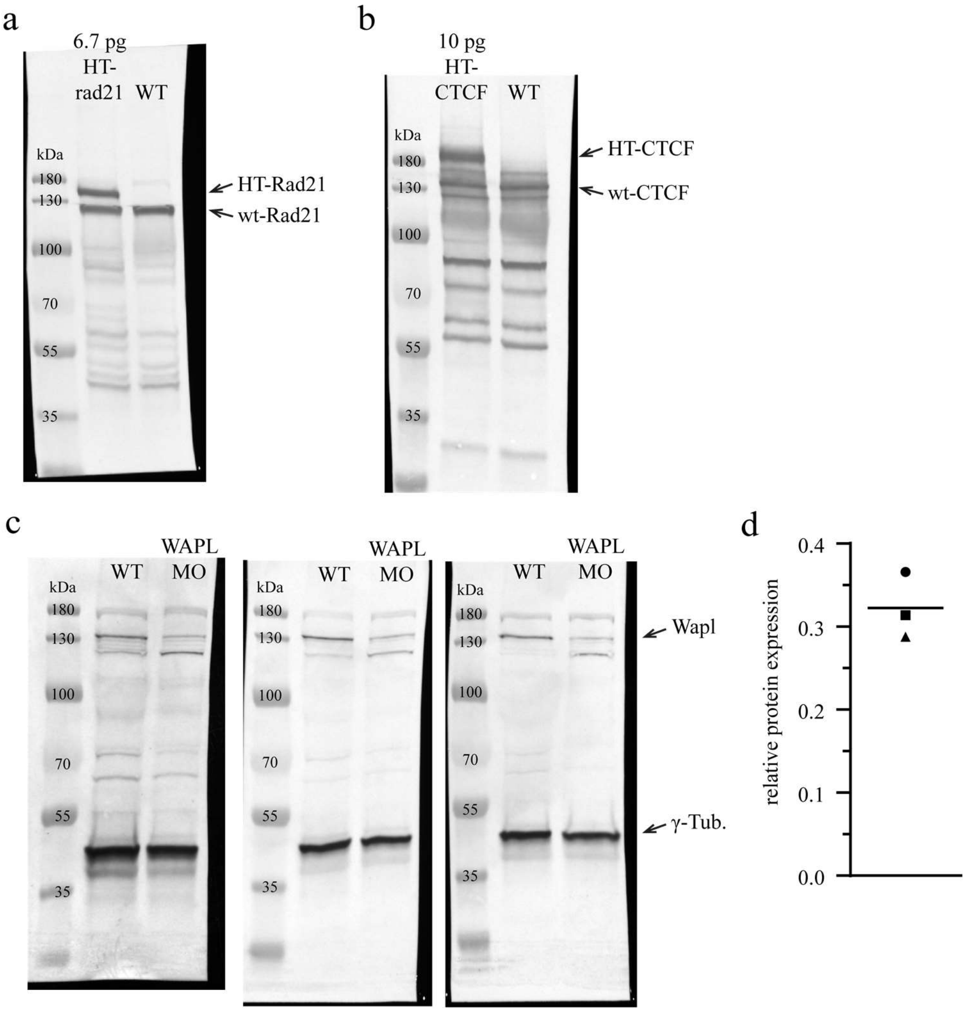

**Supplementary Figure 1. Western blots of HT-tagged proteins and after addition of Wapl-morpholino from lysates of shield-stage zebrafish embryos.** Western blots of uninjected embryos (WT) or embryos injected with **a)** 6.7 pg mRNA encoding for HT-rad21 with anti-Rad21 antibody or **b)** 10 pg mRNA encoding for HT-CTCF with anti-CTCF antibody in shield-stage. The injection amounts were increased 10-fold compared to our single-molecule measurements to enhance band clarity (see Methods). **c)** Western blots of three independent biological replicates of embryos injected with 2 ng Wapl-Morpholino (MO) with anti-Wapl and anti-γ-Tubulin (γ-Tub.) antibodies in shield-stage. **d)** Quantification of Wapl protein levels from Wapl-MO injected embryos compared to WT and loading control from Western blots shown in c). Horizontal line displays the mean. Source data are provided as a Source Data file for Supplementary Fig. 1d.

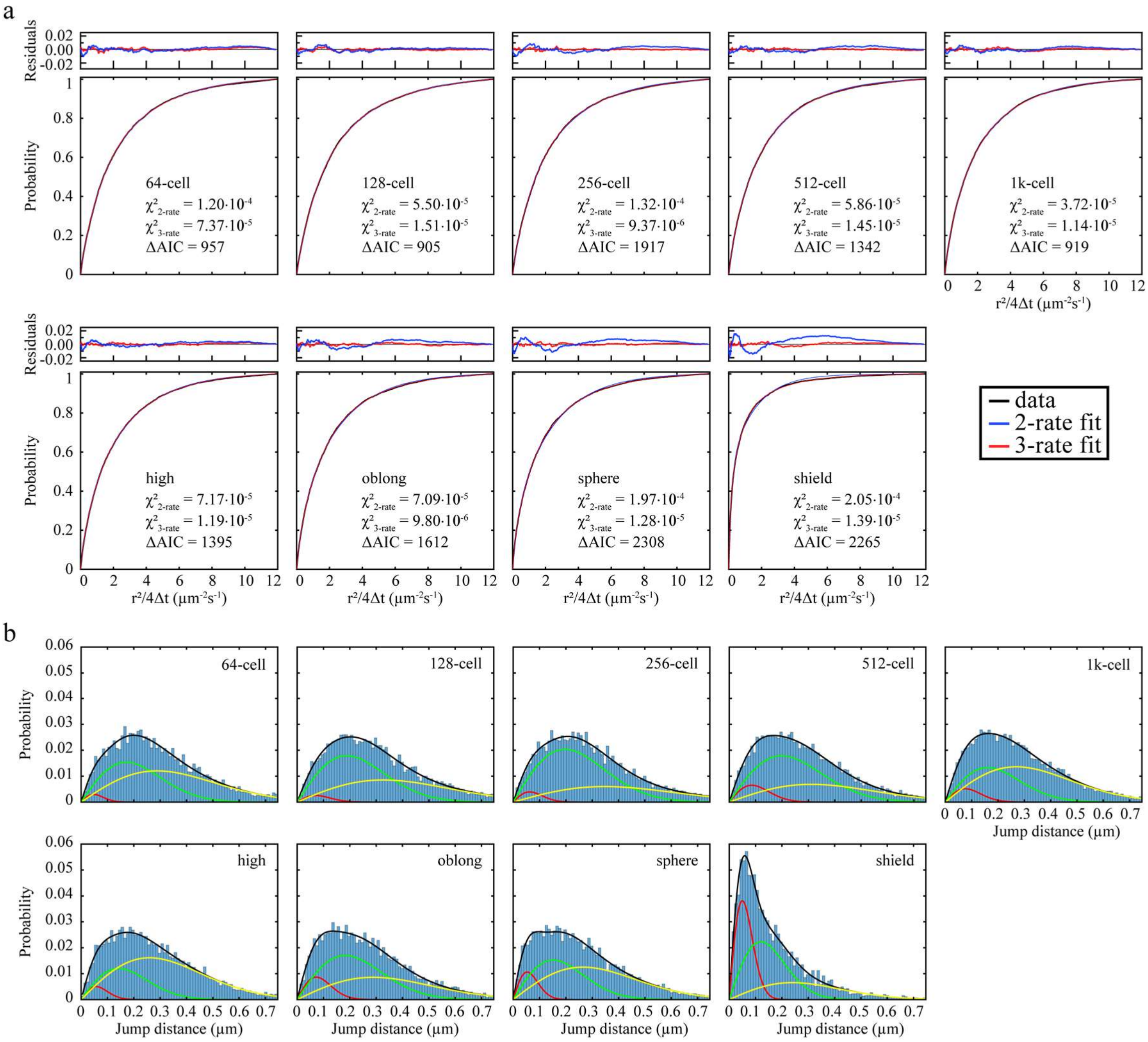

**Supplementary Figure 2. Analysis of jump distance distributions of HT-rad21 mobility data.** **a**) Cumulative distributions of jump distances (black line) taken from 11.7 ms continuous movies with fits of a 2-component (blue) and 3-component (red) diffusion model at indicated developmental stages. Values give the reduced $\chi^2$ and Akaike Information Criterion (AIC), evidencing a preferable 3-component fit. (**b**) Distributions of jump distances with a three-component diffusion model (black) and every single component (red, green, yellow) at indicated developmental stages.

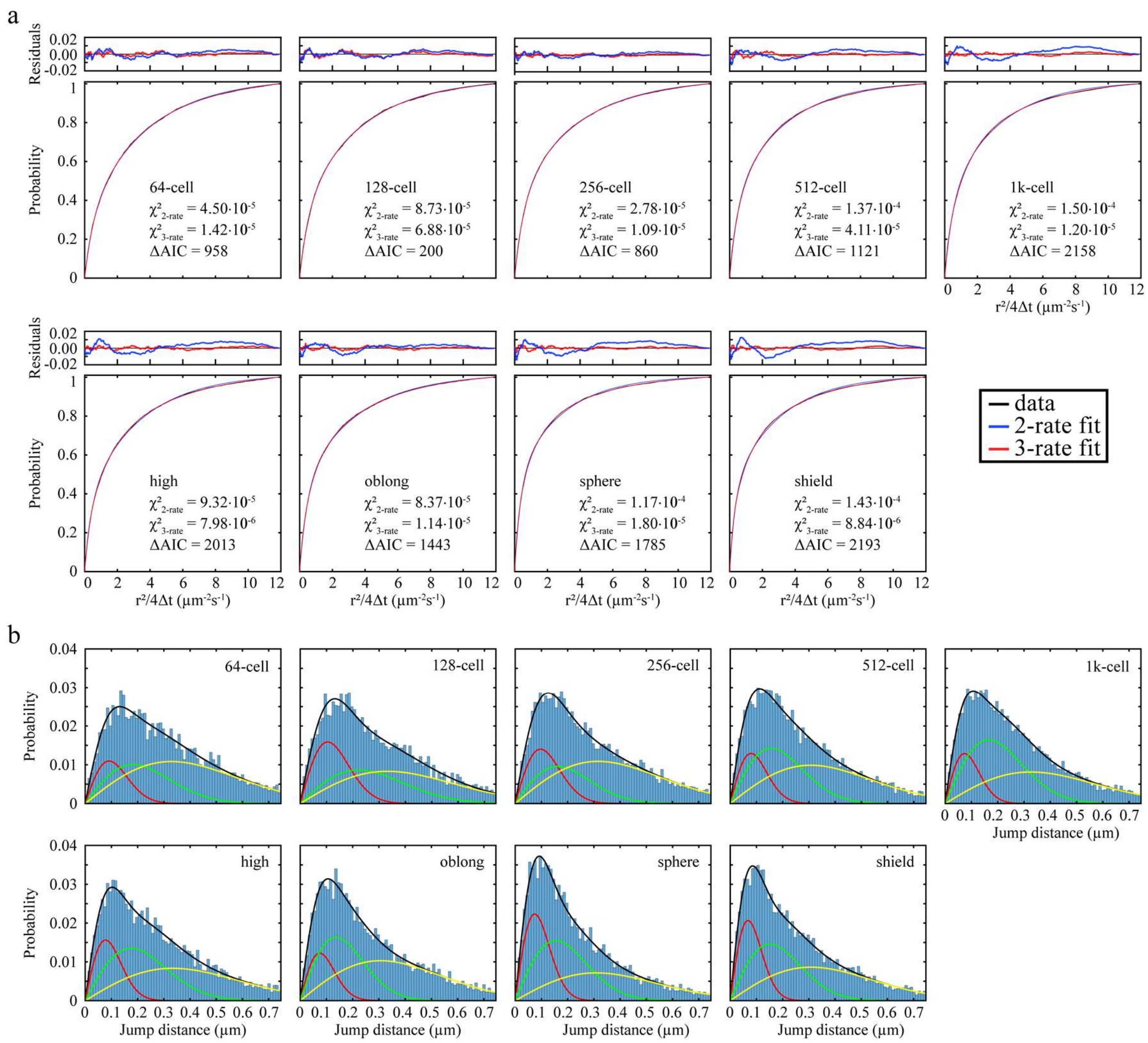

**Supplementary Figure 3. Analysis of jump distance distributions of HT-CTCF mobility data.** **a**) Cumulative distributions of jump distances (black line) taken from 11.7 ms continuous movies with fits of a 2-component (blue) and 3-component (red) diffusion model at indicated developmental stages. Values give the reduced $\chi^2$ and Akaike Information Criterion (AIC), evidencing a preferable 3-component fit. (**b**) Distribution of jump distances with a three-component diffusion model (black) and every single component (red, green, yellow) at indicated developmental stages.

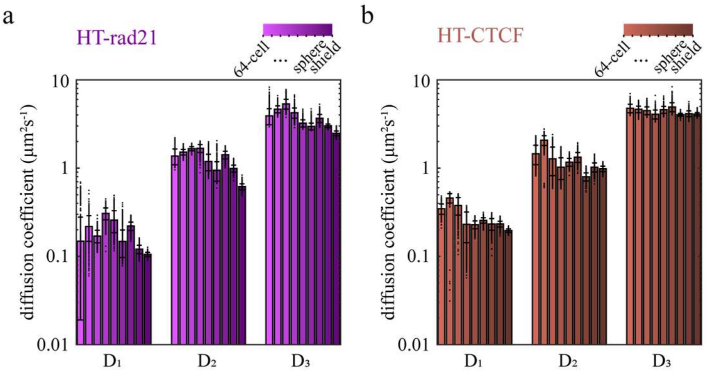

**Supplementary Figure 4. Mobility of HT-rad21 and HT-CTCF during zebrafish development.** Diffusion coefficients of **a)** HT-rad21 and **b)** HT-CTCF obtained from a three-component diffusion model fitted to jump distance distributions. Colors indicate stages of development (64-, 128-, 256-, 512-, 1k-cell, high, oblong, sphere, shield). Bars represent mean values ± s.d. from 500 resamplings using 80% of randomly selected jump distances. For amplitudes, see Fig. 1. Source data are provided as a Source Data file.

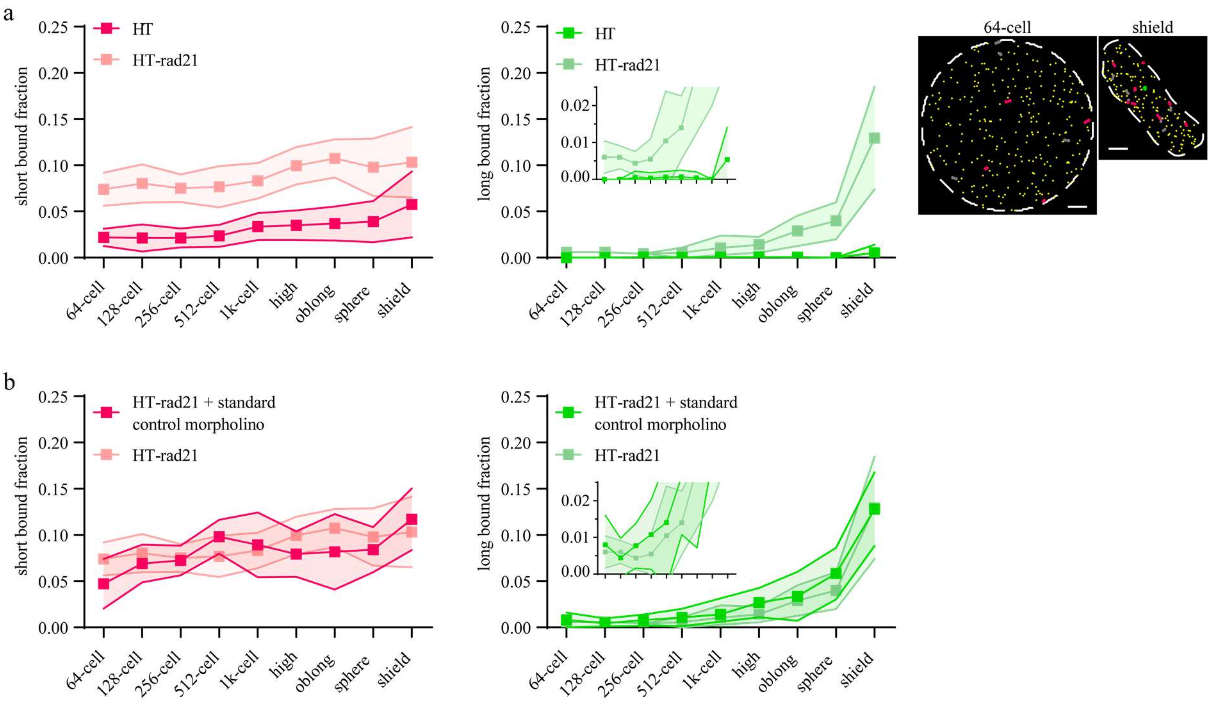

**Supplementary Figure 5. Controls for HT-rad21 recorded with interlaced time-lapse microscopy (ITM).** **a)** Left: Fractions of short and long binding events of HT control or HT-rad21 recorded with interlaced time-lapse microscopy (ITM) illumination. Right: Example nuclei from ITM movies of zebrafish embryos injected with RNA encoding for HT control. Tracks are colored according to the binding classes given in Fig. 1f. Grey tracks survived only one long dark time. Scale bar: 5 µm. **b)** Fractions of short and long binding events of HT-rad21 with coinjected standard control morpholino or HT-rad21 recorded with interlaced time-lapse microscopy (ITM) illumination. Data represent mean ± s.d. of movie-wise determined fractions. Insets show zooms in the respective graphs. Lines serve as guides to the eye. Statistics are provided in Supplementary Table 17. Source data are provided as a Source Data file.

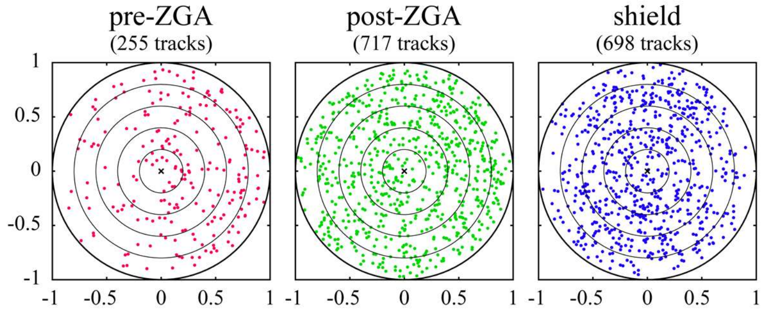

**Supplementary Figure 6. Unit circles for Center-Border Distance (CBD) analysis of HT-rad21 initial positions.** Initial positions of tracks classified as long-bound in ITM measurements (Fig 1f) and TACO measurements (Fig 4a) from pooled stages are shown on a unit circle. pre-ZGA: 64-, 128-, 256, 512-cell stages pooled; post-ZGA: high, oblong, sphere stages pooled; shield stage.

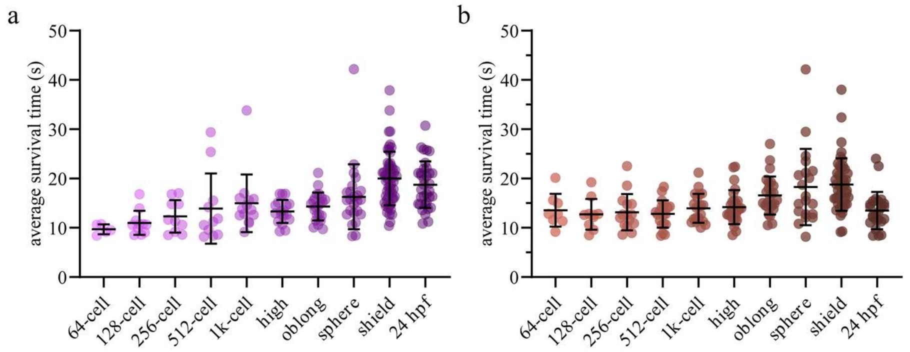

**Supplementary Figure 7. Relative increase of survival times.** Average survival times of binding events in interlaced time-lapse microscopy (ITM) that were classified as long-bound (>8.2 s) for **a**) HT-rad21 or **b)** HT-CTCF molecules. All other tracks (<8.2 s) were not considered. Data represent mean ± s.d. of movie-wise averages. Source data are provided as a Source Data file.

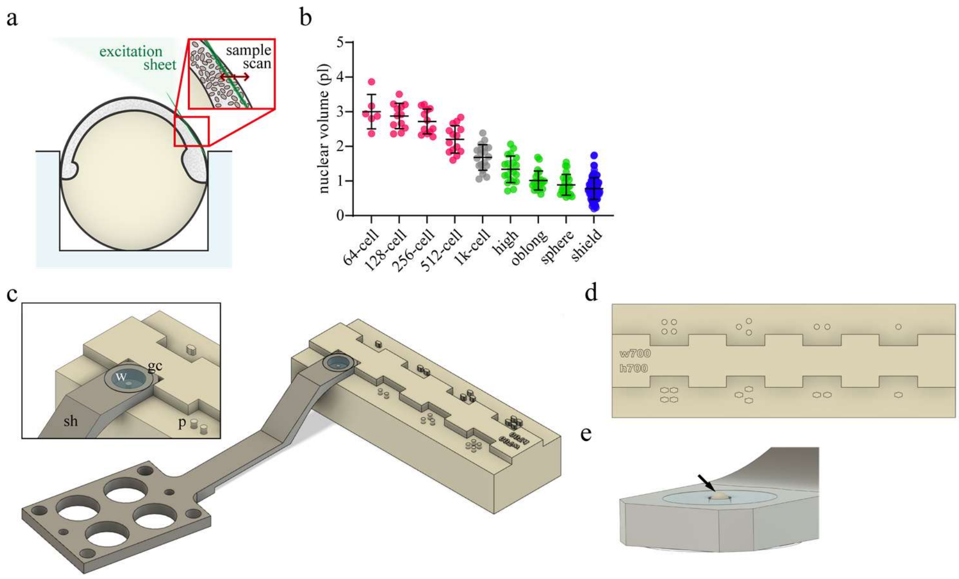

**Supplementary Figure 8. Mounting of zebrafish embryos on a lattice light-sheet microscope.** **a)** Scheme of a shield stage zebrafish embryo placed in an agarose well to scan through nuclei with a lattice light-sheet microscope. **b)** Nuclear volume at multiple stages of development. Data represent mean ± s.d. **c)** Sample holder (grey) with a 5 mm coverslip glued on top and filled with agarose. A 3D-printed stamp (brown) with pins is used to form wells in the agarose. Inset: sample holder (sh), agarose well (w), glass coverslip (gc), pins (p) **d)** Top-down view of the 3D printed stamp with multiple pin combinations for early stages up to shield stage (top row) and later stages post 24 hpf (bottom row) **e)** Zoom-in on the sample holder filled with agarose showing the animal cap of an embryo mounted in the center (arrow). The animal cap is above the agarose, allowing undisturbed excitation of fluorophores and emission of fluorescent light.

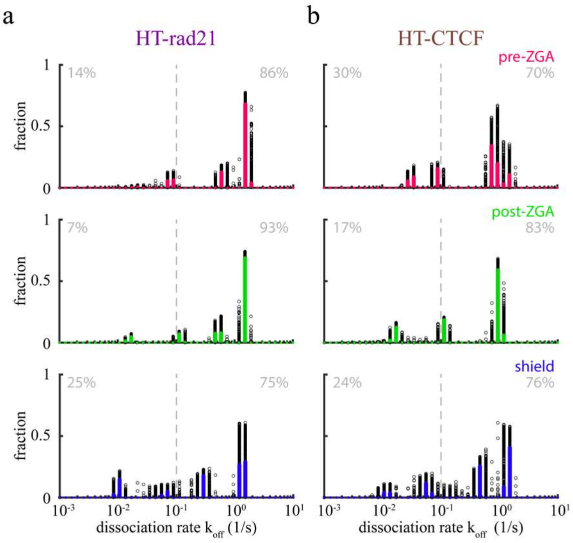

**Supplementary Figure 9. Event spectra of dissociation rates.** GRID event spectra of dissociation rates of (**a**) HT-rad21 and (**b**) HT-CTCF using all data (solid line, colored according to stages) and 500 resampling runs with randomly selected 80% of data (black spots) as an error estimation of the spectra. Grey insets: Percentages of dissociation rates larger or smaller than 0.1 $s^{-1}$ (dashed line). State spectra and residence times are provided in Fig. 3f and g. Statistics are given in Supplementary Table 14. Source data are provided as a Source Data file.

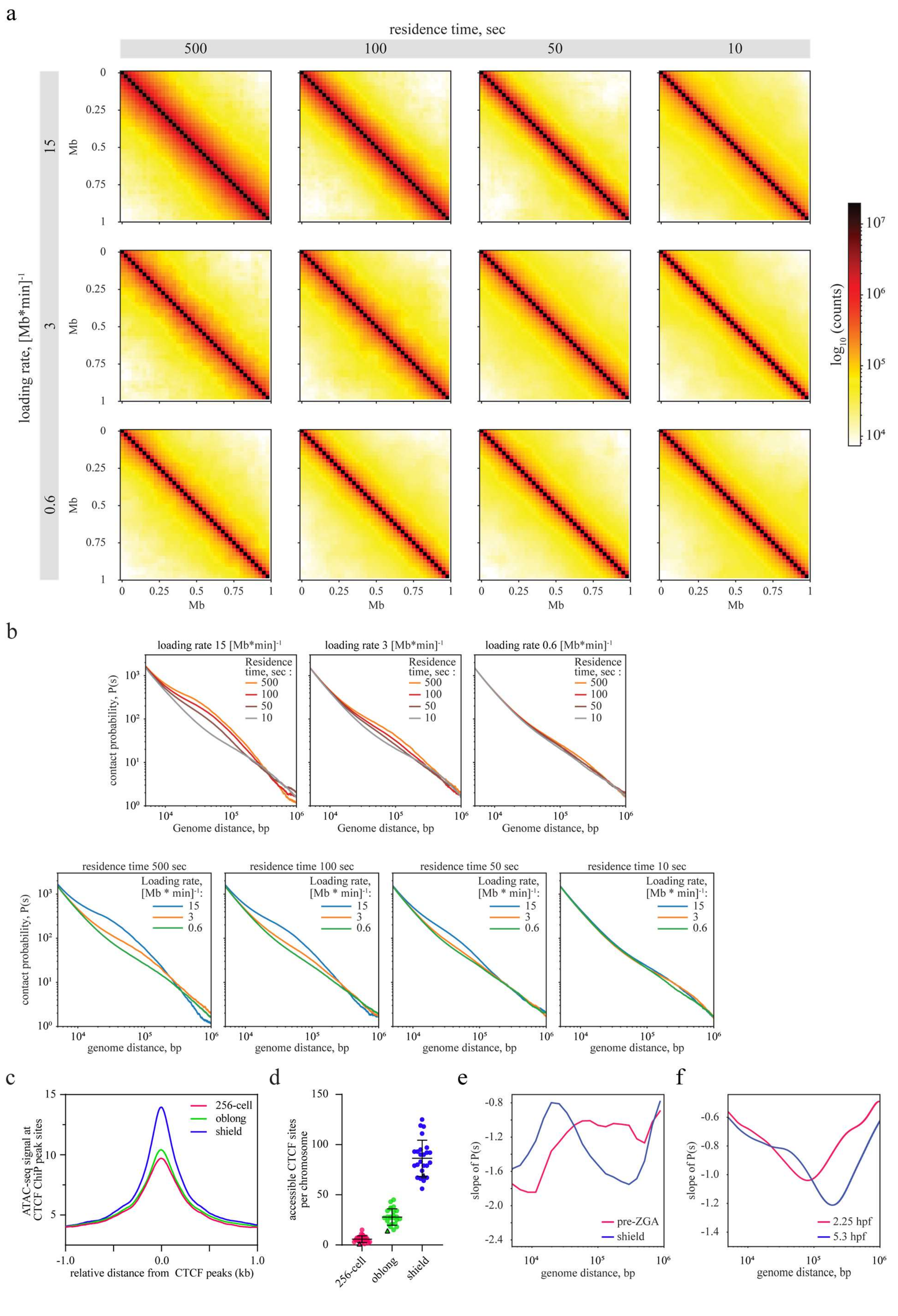

a
residence time, sec
500
100
50
10
loading rate, [Mb*min]-1
15
3
0.6
Mb
0
0.25
0.5
0.75
1
log10 (counts)
b
loading rate 15 [Mb*min]-1
loading rate 3 [Mb*min]-1
loading rate 0.6 [Mb*min]-1
Residence time, sec :
contact probability, P(s)
Genome distance, bp
residence time 500 sec
residence time 100 sec
residence time 50 sec
residence time 10 sec
Loading rate, [Mb * min]-1:
genome distance, bp
c
ATAC-seq signal at CTCF ChiP peak sites
256-cell
oblong
shield
relative distance from CTCF peaks (kb)
d
accessible CTCF sites per chromosome
256-cell
oblong
shield
e
slope of P(s)
pre-ZGA
shield
genome distance, bp
f
slope of P(s)
2.25 hpf
5.3 hpf
genome distance, bp

**Supplementary Figure 10. Sweeping of parameter space, CTCF sites and Chromatin accessibility.**
**a)** Contact maps of the polymer-chain model at 1 kb mapping. The parameter space of extruder loading rates and residence times covers 1-2 orders of magnitude. **b)** The dependence of contact probability P(s) on genomic distance for contact maps shown in panel a). **c)** ATAC-seq profile for three developmental stages at CTCF sites identified by ChIP-seq based on published data(Pálfy et al., 2019; Pérez-Rico et al., 2020) **d)** Distributions of accessible CTCF sites per chromosome identified in open chromatin regions by ATAC-seq for three developmental stages. Chr18 is highlighted with a triangle. **e)** Slopes of P(s) for pre-ZGA and shield stages from Fig. 4a. **f)** Slopes of P(s) for 2.25 hpf and 5.3 hpf from Fig. 4d. Source data are provided as a Source Data file for Suppl. Fig. 9c and d.

### Supplementary Tables

| | HT-rad21 | | | | | HT-CTCF | | | | |
|---|---|---|---|---|---|---|---|---|---|---|
| | Days | Embryo | Movie | Tracks | Jumps | Days | Embryo | Movie | Tracks | Jumps |
| 64-cell | 4 | 7 | 10 | 1474 | 8432 | 2 | 4 | 9 | 1654 | 8992 |
| 128-cell | 4 | 9 | 13 | 1828 | 9744 | 2 | 6 | 10 | 1455 | 7675 |
| 256-cell | 4 | 8 | 13 | 1939 | 12222 | 2 | 7 | 14 | 1859 | 10060 |
| 512-cell | 4 | 12 | 16 | 1697 | 11517 | 2 | 7 | 13 | 1616 | 10406 |
| 1k-cell | 4 | 11 | 21 | 1888 | 12932 | 2 | 8 | 15 | 1900 | 12585 |
| high | 4 | 9 | 14 | 1376 | 10613 | 2 | 8 | 16 | 1773 | 11856 |
| oblong | 4 | 6 | 11 | 1123 | 9049 | 2 | 7 | 15 | 1507 | 10915 |
| sphere | 4 | 9 | 21 | 1391 | 12035 | 2 | 7 | 20 | 1870 | 15339 |
| shield | 3 | 12 | 29 | 922 | 9637 | 2 | 9 | 35 | 2422 | 17525 |

**Supplementary Table 1.** Statistics of 11.7 ms continuous movies recorded for HT-rad21 and HT-CTCF. Related to Fig. 1d,e.

| | long bound fraction | short bound fraction |
|---|---|---|
| 64-cell | 0.0030 | 0.1079 |
| 128-cell | 0.0002 | 0.0409 |
| 256-cell | 0.0032 | 0.0409 |
| 512-cell | 0.0388 | 0.7267 |
| 1k-cell | 0.0051 | 0.2864 |
| high | <0.0001 | 0.1968 |
| oblong | <0.0001 | 0.1762 |
| sphere | <0.0001 | 0.4819 |
| shield | <0.0001 | 0.1645 |
| 24 hpf | <0,0001 | 0.2345 |

**Supplementary Table 2.** Comparison of bound fractions between HT-rad21 and HT-rad21-3x. P-values from multiple Mann-Whitney tests on interlaced time-lapse microscopy (ITM) data shown in Fig. 1h.

| | long bound fraction | short bound fraction |
|---|---|---|
| 64-cell | 0.0009 | 0.7802 |
| 128-cell | 0.0002 | 0.6522 |
| 256-cell | <0.0001 | 0.0105 |
| 512-cell | <0.0001 | 0.0361 |
| 1k-cell | <0.0001 | 0.0220 |
| high | <0.0001 | 0.2041 |
| oblong | <0.0001 | 0.0093 |
| sphere | <0.0001 | 0.0133 |
| shield | <0.0001 | 0.0008 |
| 24 hpf | <0.0001 | 0.6419 |

**Supplementary Table 3.** Comparison of bound fractions between HT-CTCF and HT-CTCF-ΔZF4-7. P-values from multiple Mann-Whitney tests on interlaced time-lapse microscopy (ITM) data shown in Fig. 1g.

| | **long-bound fractions** | | | | | | | | |
|---|---|---|---|---|---|---|---|---|---|
| | 128-cell | 256-cell | 512-cell | 1k-cell | high | oblong | sphere | shield | 24 hpf |
| 64-cell | >0.9999 | >0.9999 | >0.9999 | >0.9999 | >0.9999 | >0.9999 | 0.8898 | 0.0001 | <0.0001 |
| 128-cell | | >0.9999 | >0.9999 | >0.9999 | >0.9999 | 0.8844 | 0.1341 | <0.0001 | <0.0001 |
| 256-cell | | | >0.9999 | >0.9999 | >0.9999 | 0.2909 | 0.0343 | <0.0001 | <0.0001 |
| 512-cell | | | | >0.9999 | >0.9999 | 0.3452 | 0.0373 | <0.0001 | <0.0001 |
| 1k-cell | | | | | >0.9999 | 0.9325 | 0.1024 | <0.0001 | <0.0001 |
| high | | | | | | >0.9999 | >0.9999 | <0.0001 | <0.0001 |
| oblong | | | | | | | >0.9999 | 0.0002 | <0.0001 |
| sphere | | | | | | | | 0.0044 | 0.0003 |
| shield | | | | | | | | | >0.9999 |

| | **short-bound fractions** | | | | | | | | |
|---|---|---|---|---|---|---|---|---|---|
| | 128-cell | 256-cell | 512-cell | 1k-cell | high | oblong | sphere | shield | 24 hpf |
| 64-cell | >0.9999 | >0.9999 | >0.9999 | >0.9999 | >0.9999 | 0.2965 | >0.9999 | 0.8475 | 0.0590 |
| 128-cell | | >0.9999 | >0.9999 | >0.9999 | >0.9999 | 0.4666 | >0.9999 | >0.9999 | 0.0517 |
| 256-cell | | | >0.9999 | >0.9999 | 0.4719 | 0.0236 | >0.9999 | 0.0732 | 0.0011 |
| 512-cell | | | | >0.9999 | >0.9999 | 0.0592 | >0.9999 | 0.1878 | 0.0029 |
| 1k-cell | | | | | >0.9999 | 0.3497 | >0.9999 | >0.9999 | 0.0202 |
| high | | | | | | >0.9999 | >0.9999 | >0.9999 | >0.9999 |
| oblong | | | | | | | >0.9999 | >0.9999 | >0.9999 |
| sphere | | | | | | | | >0.9999 | 0.7912 |
| shield | | | | | | | | | >0.9999 |

**Supplementary Table 4.** Comparison of bound fractions of HT-rad21. P-values from Kruskal-Wallis multiple comparison tests on interlaced time-lapse microscopy (ITM) data shown in Fig. 1h.

| | **long-bound fractions** | | | | | | | | |
|---|---|---|---|---|---|---|---|---|---|
| | 128-cell | 256-cell | 512-cell | 1k-cell | high | oblong | sphere | shield | 24 hpf |
| 64-cell | >0.9999 | >0.9999 | >0.9999 | >0.9999 | 0.1876 | 0.0005 | 0.0019 | 0.0007 | 0.0008 |
| 128-cell | | >0.9999 | >0.9999 | 0.8534 | 0.0371 | <0.0001 | 0.0002 | <0.0001 | <0.0001 |
| 256-cell | | | >0.9999 | >0.9999 | 0.2592 | 0.0002 | 0.0012 | 0.0002 | 0.0003 |
| 512-cell | | | | >0.9999 | >0.9999 | 0.0121 | 0.0490 | 0.0133 | 0.0195 |
| 1k-cell | | | | | >0.9999 | 0.0804 | 0.2552 | 0.1060 | 0.1213 |
| high | | | | | | >0.9999 | >0.9999 | >0.9999 | >0.9999 |
| oblong | | | | | | | >0.9999 | >0.9999 | >0.9999 |
| sphere | | | | | | | | >0.9999 | >0.9999 |
| shield | | | | | | | | | >0.9999 |

| | **short-bound fractions** | | | | | | | | |
|---|---|---|---|---|---|---|---|---|---|
| | 128-cell | 256-cell | 512-cell | 1k-cell | high | oblong | sphere | shield | 24 hpf |
| 64-cell | >0.9999 | >0.9999 | 0.5778 | 0.1195 | 0.1444 | 0.3779 | 0.0147 | 0.0005 | 0.0009 |
| 128-cell | | >0.9999 | >0.9999 | 0.9423 | >0.9999 | >0.9999 | 0.1467 | 0.0069 | 0.0117 |
| 256-cell | | | >0.9999 | >0.9999 | >0.9999 | >0.9999 | >0.9999 | 0.4751 | 0.5842 |
| 512-cell | | | | >0.9999 | >0.9999 | >0.9999 | >0.9999 | >0.9999 | >0.9999 |
| 1k-cell | | | | | >0.9999 | >0.9999 | >0.9999 | >0.9999 | >0.9999 |
| high | | | | | | >0.9999 | >0.9999 | >0.9999 | >0.9999 |
| oblong | | | | | | | >0.9999 | 0.8510 | >0.9999 |
| sphere | | | | | | | | >0.9999 | >0.9999 |
| shield | | | | | | | | | >0.9999 |

**Supplementary Table 5.** Comparison of bound fractions of HT-CTCF. P-values from Kruskal-Wallis multiple comparison tests on interlaced time-lapse microscopy (ITM) data shown in Fig. 1g.

| | HT-rad21 | | | | HT-rad21-3x mutant | | | |
|---|---|---|---|---|---|---|---|---|
| | Days | Embryos | Movies | Mean number of all events per movie | Days | Embryos | Movies | Mean number of all events per movie |
| 64-cell | 4 | 6 | 6 | 752 | 2 | 6 | 8 | 427 |
| 128-cell | 4 | 8 | 12 | 676 | 2 | 6 | 9 | 541 |
| 256-cell | 4 | 8 | 12 | 573 | 2 | 5 | 9 | 421 |
| 512-cell | 4 | 9 | 14 | 506 | 2 | 6 | 11 | 366 |
| 1k-cell | 4 | 10 | 19 | 429 | 2 | 6 | 10 | 372 |
| high | 4 | 11 | 20 | 399 | 2 | 6 | 11 | 293 |
| oblong | 4 | 11 | 22 | 331 | 2 | 6 | 10 | 230 |
| sphere | 4 | 9 | 24 | 245 | 2 | 6 | 14 | 149 |
| shield | 2 | 9 | 56 | 68 | 2 | 8 | 38 | 40 |
| 24 hpf | 2 | 11 | 34 | 45 | 2 | 6 | 14 | 105 |

**Supplementary Table 6.** Statistics of interlaced time-lapse microscopy (ITM) movies recorded for HT-rad21 and HT-rad21-3x. Related to ITM data in Fig. 1h.

| | HT-CTCF | | | | HT-CTCF-ΔZF4-7 mutant | | | |
|---|---|---|---|---|---|---|---|---|
| | Days | Embryos | Movies | Mean number of all events per movie | Days | Embryos | Movies | Mean number of all events per movie |
| 64-cell | 3 | 5 | 9 | 284 | 2 | 6 | 10 | 718 |
| 128-cell | 4 | 6 | 11 | 264 | 2 | 7 | 11 | 769 |
| 256-cell | 4 | 7 | 16 | 241 | 2 | 7 | 11 | 796 |
| 512-cell | 5 | 10 | 20 | 245 | 2 | 7 | 14 | 813 |
| 1k-cell | 5 | 8 | 21 | 256 | 2 | 6 | 11 | 673 |
| high | 5 | 10 | 26 | 211 | 2 | 6 | 11 | 666 |
| oblong | 5 | 9 | 25 | 175 | 2 | 6 | 13 | 555 |
| sphere | 3 | 6 | 20 | 133 | 2 | 6 | 12 | 452 |
| shield | 2 | 11 | 46 | 195 | 2 | 9 | 50 | 324 |
| 24 hpf | 2 | 7 | 24 | 92 | 2 | 7 | 23 | 184 |

**Supplementary Table 7.** Statistics of interlaced time-lapse microscopy (ITM) movies recorded for HT-CTCF and HT-CTCF-ΔZF4-7. Related to ITM data in Fig. 1g.

| HT-rad21+ | **short-bound** | **long-bound** |
|---|---|---|
| ctcf-MO | >0,9999 | <0,0001 |
| nipbl-MO | >0,9999 | 0,1371 |
| wapl-MO | >0,9999 | 0,0066 |
| α-Amanitin | 0,0005 | 0,0001 |
| Triptolide | 0,0021 | 0,3718 |

**Supplementary Table 8.** Comparison of bound fractions of HT-rad21 wild-type vs. morpholino addition (MO) or RNA-Polymerase inhibitors for interlaced time-lapse microscopy (ITM) data in shield-stage. P-values from Kruskal-Wallis multiple comparison tests on data shown in Fig. 1i.

| (shield stage) HT-rad21+ | Days | Embryos | Movies | Mean number of all events per movie |
|---|---|---|---|---|
| ctcf-MO | 2 | 5 | 41 | 122 |
| nipbl-MO | 2 | 9 | 54 | 94 |
| wapl-MO | 3 | 11 | 27 | 144 |
| α-Amanitin | 2 | 4 | 27 | 126 |
| Triptolide | 2 | 9 | 21 | 96 |

**Supplementary Table 9.** Statistics of bound fractions of HT-rad21 wild-type vs. morpholino addition (MO) or RNA-Polymerase Inhibitors for interlaced time-lapse microscopy (ITM) measurements. Related to Fig. 1i.

| | **short-bound** | | **long-bound** | | |
|---|---|---|---|---|---|
| | post-ZGA | shield | post-ZGA | shield | shield + nibpl-MO |
| pre-ZGA | 0.0591 | <0,0001 | <0,0001 | <0,0001 | <0,0001 |
| post-ZGA | | <0,0001 | | <0,0001 | >0,9999 |
| shield | | | | | <0,0001 |

**Supplementary Table 10.** Comparison of stages and morpholino addition (MO) of HT-rad21 for time-lapse alternated with continuous intervals (TACO) measurements. P-values from Kruskal-Wallis multiple comparison tests on data shown in Fig. 4b.

| | | Days | Embryos | Movies | Tracks |
|---|---|---|---|---|---|
| short bound | pre-ZGA | 3 | 8 | 36 | 200 |
| | post-ZGA | 3 | 7 | 36 | 221 |
| | shield | 4 | 15 | 43 | 135 |
| long bound | pre-ZGA | 4 | 11 | 33 | 80 |
| | post-ZGA | 4 | 11 | 41 | 173 |
| | shield | 4 | 15 | 42 | 248 |
| | shield +nibpl-MO | 3 | 13 | 41 | 412 |

**Supplementary Table 11.** Statistics of time-lapse alternated with continuous intervals (TACO) movies recorded for HT-rad21. Related to Fig. 3b.

| | | **Cluster** | | | |
|---|---|---|---|---|---|
| | | #1 (longest) | #2 | #3 | #4 (shortest) |
| pre-ZGA | residence time of cluster ± s.d. of resampling (s) | 52.08 ( ± 5.7) | 12.74 ( ± 0.62) | 1.66 ( ± 0.14) | 0.65 ( ± 0.03) |
| | fraction of cluster ± s.d. of resampling | 25.3 ( ± 3.3) | 52.2 ( ± 3) | 7.39 ( ± 1.2) | 15.1 ( ± 0.89) |
| post-ZGA | residence time of cluster ± s.d. of resampling (s) | 62.89 ( ± 1.9) | 8.62 ( ± 0.59) | 1.88 ( ± 0.17) | 0.67 ( ± 0.01) |
| | fraction of cluster ± s.d. of resampling | 74.5 ( ± 1.1) | 12.2 ( ± 0.75) | 5.33 ( ± 0.63) | 8.02 ( ± 0.45) |
| shield | residence time of cluster ± s.d. of resampling (s) | 99.01 ( ± 5.69) | 15.92 ( ± 3.55) | 3.53 ( ± 0.42) | 0.75 ( ± 0.06) |
| | fraction of cluster ± s.d. of resampling | 88.6 ( ± 1.9) | 6.05 ( ± 1.7) | 3.27 ( ± 0.58) | 2.12 ( ± 0.17) |

**Supplementary Table 12.** Residence times of HT-rad21. Residence times obtained from dissociation rate clusters of GRID state spectra shown in Fig. 2f.

| | | Cluster | | |
|---|---|---|---|---|
| | | #1 (longest) | #2 | #3 (shortest) |
| pre-ZGA | residence time of cluster ± s.d. of resampling (s) | 32.57 ( ± 1.8) | 11.52 ( ± 0.94) | 1.13 ( ± 0.04) |
| | fraction of cluster ± s.d. of resampling | 63.4 ( ± 4.1) | 26 ( ± 4) | 10.6 ( ± 0.73) |
| post-ZGA | residence time of cluster ± s.d. of resampling (s) | 61.35 ( ± 2.63) | 9.01 ( ± 0.48) | 1.04 ( ± 0.04) |
| | fraction of cluster ± s.d. of resampling | 79.1 ( ± 1.2) | 14.9 ( ± 0.94) | 5.94 ( ± 0.39) |
| shield | residence time of cluster ± s.d. of resampling (s) | 88.5 ( ± 10.96) | 17.33 ( ± 1.83) | 1.19 ( ± 0.07) |
| | fraction of cluster ± s.d. of resampling | 65.3 ( ± 4.2) | 26.6 ( ± 4) | 8.09 ( ± 1.1) |

**Supplementary Table 13.** Residence times of HT-CTCF. Residence times obtained from dissociation rate clusters of GRID state spectra shown in Fig. 2g.

| | | Days | Embryos | Movies | Tracks | Tracks per movie |
|---|---|---|---|---|---|---|
| HT-rad21 502ms | preZGA | 3 | 10 | 62 | 1582 | 25.5 |
| | postZGA | 3 | 9 | 59 | 2580 | 43.7 |
| | shield | 3 | 10 | 68 | 1561 | 23.0 |
| HT-rad21 4.5s | preZGA | 4 | 9 | 16 | 769 | 48.1 |
| | postZGA | 4 | 12 | 23 | 1272 | 55.3 |
| | shield | 4 | 10 | 22 | 517 | 23.5 |
| HT-CTCF 502ms | preZGA | 4 | 11 | 33 | 2334 | 70.7 |
| | postZGA | 4 | 11 | 24 | 1486 | 61.9 |
| | shield | 4 | 15 | 18 | 1278 | 71.0 |
| HT-CTCF 4.5s | preZGA | 4 | 10 | 16 | 1734 | 108.4 |
| | postZGA | 4 | 10 | 22 | 1576 | 71.6 |
| | shield | 4 | 8 | 11 | 774 | 70.4 |

**Supplementary Table 14.** Statistics of time-lapse microscopy movies recorded for HT-rad21 and HT-CTCF. Related to Fig. 2d-g.

| | $k_{off,u}$ | $A^e_s$ | $A^s_u$ | $f_b$ |
|---|---|---|---|---|
| pre-ZGA | 0.835 | 0.30 | 0.12 | 0.20 |
| post-ZGA | 0.386 | 0.17 | 0.19 | 0.21 |
| shield | 0.792 | 0.24 | 0.09 | 0.23 |

**Supplementary Table 15.** Measured parameters used to calculate the search time $\tau_{search}$ for a single HT-CTCF molecule. Related to Fig. 3i and Methods.

| | $k_{off,s}$ | $A^s_s$ | $f_b$ |
|---|---|---|---|
| pre-ZGA | 0.0591 | 0.77 | 0.05 |
| post-ZGA | 0.0165 | 0.75 | 0.08 |
| shield | 0.0132 | 0.95 | 0.31 |

**Supplementary Table 16.** Measured parameters used to calculate the search time $\tau_{search}$ for a single HT-rad21 molecule. Related to Fig. 3i and Methods.

| | HT only | | | | MO standard control | | | |
|---|---|---|---|---|---|---|---|---|
| | Days | Embryos | Movies | Mean number of all events per movie | Days | Embryos | Movies | Mean number of all events per movie |
| 64-cell | 3 | 5 | 10 | 216 | 3 | 5 | 7 | 202 |
| 128-cell | 3 | 7 | 15 | 193 | 3 | 6 | 11 | 265 |
| 256-cell | 3 | 7 | 16 | 179 | 3 | 7 | 17 | 318 |
| 512-cell | 3 | 7 | 19 | 194 | 3 | 6 | 13 | 332 |
| 1k-cell | 3 | 6 | 16 | 165 | 2 | 5 | 11 | 286 |
| high | 3 | 7 | 16 | 164 | 2 | 4 | 11 | 216 |
| oblong | 3 | 6 | 13 | 148 | 3 | 5 | 13 | 192 |
| sphere | 3 | 4 | 11 | 143 | 3 | 5 | 8 | 138 |
| shield | 3 | 10 | 34 | 89 | 3 | 9 | 40 | 142 |

**Supplementary Table 17.** Statistics of interlaced time-lapse microscopy (ITM) movies recorded for HT-control and HT-rad21 with coinjected standard control morpholino. Related to Supplementary Fig. 5.

| | |
|---|---|
| CTCF fwd PacI | GAACTTTAATTAATATGGAAGGGGGACCGAC |
| CTCF rev AscI | GAACTGGCGCGCCTCACCGGTCCATCATGCTAAG |
| Rad21a fwd PacI | GAACTTTAATTAAGATGTTTTACGCCCACTTCGTC |
| Rad21a rev AscI | GAACTGGCGCGCCCTATACAATGTGGAAGCGTGGT |
| Rad21a 3x fwd Q5SDM | GCAGCAGGCCATCGACCTGACGAAGACCGAGCCCTACAGTGAC |
| Rad21a 3x rev Q5SDM | TTCTTCAGCACCCGGAAGCTGTAACGCTTGGCCGCAGCCTGTTT |
| CTCF ΔZF47 fwd Q5SDM: | AGAAAGTGCCGTTACTGTG |
| CTCF ΔZF47 rev Q5SDM: | CGGTTTCTCATGAGTGTG |

**Supplementary Table 18.** Primers for cloning of HT constructs and Q5 Site-Directed Mutagenesis (Q5SDM).

| | |
|---|---|
| MO wapla ATG | TTATCGATCCTCCGTCTTTCCTCTG |
| MO waplb ATG | CATCTGTCTCATATTCTCTGGTTGG |

**Supplementary Table 19.** Wapl morpholino (MO) sequences complementary to the target sequence.

| | Frame cycle time (ms) | Excitation sequence |
|---|---|---|
| 11.7 ms continuous | 11.7 | 117 ms/10f 488, 1.17 s/100f 561 |
| Interlaced time-lapse microscopy (ITM) | 11.7 | 11x (2x (11.7 ms/1f 561, 117 ms/10f 488, 70.2 ms dark), 1.80 s dark, 117 ms/10f 488, 1.87 s dark) |
| 0.5 s continuous time-lapse | 501.7 | 501.7 ms/1f 488, 60.20 s/120f 561, 501.7 ms/1f 488 |
| 4.5 s time-lapse | 501.7 | 501.7 ms/1f 561, 501.7 ms/1f 488, 3.51 s dark |
| Time-lapse alternated with continuous intervals (TACO) short | 11.7 | 11x (11.7 ms/1f 561, 117 ms/10f 488, 70.2 ms dark, 117 ms/10f 561, 117 ms/10f 488, 3.76 s dark) |
| TACO long | 11.7 | 3.6x (2x (11.7 ms /1f 561, 117 ms/10f 488, 4.06 s dark), 117 ms/10f 561, 1.92 s dark, 117 ms/10f 488, 2.04 s dark) |

**Supplementary Table 20.** Illumination scheme names, frame cycle times (= exposure times + 1.7 ms readout time), and excitation sequences are given in time and frame counts (f), followed by the excitation laser wavelengths (488 nm, 561 nm).

| | Construct | Threshold factor | Tracking radius (µm) | Min track length | Gap frames | Min. track length before gap frame |
|---|---|---|---|---|---|---|
| 11.7 ms continuous | HT-rad21, HT-CTCF | 1 | 0.747 | 2 | 1 | 2 |
| Interlaced time-lapse microscopy (ITM) | HT-rad21, HT-rad21-3x, HT-CTCF, HT-CTCF-ΔZF4-7 | 1 | 0.498 | 2 | 0 | 0 |
| 0.5 s continuous time-lapse | HT-rad21, HT-CTCF | 1 | 0.370 | 3 | 1 | 2 |
| 4.5 s time-lapse | HT-rad21, HT-CTCF | 1 | 0.907 | 3 | 1 | 2 |
| Time-lapse alternated with continuous intervals (TACO) unspecific | HT-rad21 | 1 | 0.747 | 2 | 1 | 2 |
| TACO specific | HT-rad21 | 1 | 0.747 | 2 | 1 | 2 |

**Supplementary Table 21.** Tracking parameters used in TrackIt for our illumination schemes, all using the nearest neighbor algorithm.

| >GCCWGCAGGGGGCGCTGSDG CTCF_zebrafish 3.643980 -616.688268 0 T:1993.0(42.57%),B:7136.0(20.02%),P:1e-267 | | | |
|---|---|---|---|
| 0.125 | 0.078 | 0.550 | 0.247 |
| 0.047 | 0.576 | 0.309 | 0.068 |
| 0.046 | 0.908 | 0.015 | 0.031 |
| 0.453 | 0.046 | 0.171 | 0.330 |
| 0.015 | 0.108 | 0.876 | 0.001 |
| 0.061 | 0.892 | 0.046 | 0.001 |
| 0.953 | 0.001 | 0.015 | 0.031 |
| 0.001 | 0.001 | 0.997 | 0.001 |
| 0.123 | 0.001 | 0.861 | 0.015 |
| 0.078 | 0.123 | 0.732 | 0.067 |
| 0.001 | 0.001 | 0.997 | 0.001 |
| 0.015 | 0.001 | 0.969 | 0.015 |
| 0.001 | 0.997 | 0.001 | 0.001 |
| 0.202 | 0.015 | 0.782 | 0.001 |
| 0.001 | 0.860 | 0.124 | 0.015 |
| 0.217 | 0.281 | 0.062 | 0.440 |
| 0.092 | 0.092 | 0.815 | 0.001 |
| 0.123 | 0.396 | 0.357 | 0.124 |
| 0.283 | 0.157 | 0.219 | 0.341 |
| 0.062 | 0.219 | 0.611 | 0.108 |

**Supplementary Table 22.** CTCF motif for CTCF orientation analysis. See Methods. Motif from a published dataset (Pérez-Rico et al., 2020) with GEO accession number GSE133437.

#### Supplementary Movie Legends

**File name: Supplementary Movie 1**
Description: Example movie of HT-rad21 mobility in 64-cell stage zebrafish embryo. Left: single molecule movie of HT-rad21 molecules with continuous illumination at 11.7 ms frame cycle time (see Fig. 1c). Right: signal of the Lap2β nuclear membrane marker. Scale bar: 5 µm.

**File name: Supplementary Movie 2**
Description: Example movie of HT-rad21 mobility in shield stage zebrafish embryo**.** Left: single molecule movie of HT-rad21 molecules with continuous illumination at 11.7 ms frame cycle time (see Fig. 1c). Right: signal of the Lap2β nuclear membrane marker. Scale bar: 5 µm.

**File name: Supplementary Movie 3**
Description: Example movie of HT-rad21 binding classes in 64-cell stage zebrafish embryos. Left: single molecule movie of HT-rad21 molecules recorded with interlaced time-lapse microscopy (ITM) illumination (see Fig. 1f). Right: signal of the Lap2β nuclear membrane marker. Tracks are colored according to binding classes: long (green), intermediate (grey), and short (red). Scale bar: 5 µm.

**File name: Supplementary Movie 4**
Description: Example movie of HT-rad21 binding classes in 24hpf zebrafish embryos. Left: single molecule movie of HT-rad21 molecules recorded with interlaced time-lapse microscopy (ITM) illumination (see Fig. 1f). Right: signal of the Lap2β nuclear membrane marker. Tracks are colored according to binding classes: long (green), intermediate (grey), and short (red). Scale bar: 5 µm.

**File name: Supplementary Movie 5**
Description: Example movie of long-bound HT- rad21 mobility in 64-cell stage zebrafish embryos. Left: single molecule movie of HT-rad21 molecules recorded with the long time-lapse alternated with continuous intervals (TACO) illumination (see Fig. 4a). Right: signal of the Lap2β nuclear membrane marker. Only the first continuous illumination period of tracks that meet the long detection definition are shown. Scale bar: 5 µm.

**File name: Supplementary Movie 6**
Description: Example movie of long-bound HT- rad21 mobility in shield stage zebrafish embryos. Left: single molecule movie of HT-rad21 molecules recorded with the long time-lapse alternated with continuous intervals (TACO) illumination (see Fig. 4a). Right: signal of the Lap2β nuclear membrane marker. Only the first continuous illumination period of tracks that meet the long detection definition are shown. Scale bar: 5 µm.

**File name: Supplementary Movie 7**
Description: Example movie of short-bound HT- rad21 mobility in 64-cell stage zebrafish embryos. Left: single molecule movie of HT-rad21 molecules recorded with the short time-lapse alternated with continuous intervals (TACO) illumination (see Fig. 4a). Right: signal of the Lap2β nuclear membrane marker. Only tracks that meet the short detection definition are shown. Scale bar: 5 µm.

**File name: Supplementary Movie 8**
Description: Example movie of short-bound HT- rad21 mobility in shield stage zebrafish embryos. Left: single molecule movie of HT-rad21 molecules recorded with the short time-lapse alternated with continuous intervals (TACO) illumination (see Fig. 4a). Right: signal of the Lap2β nuclear membrane marker. Only tracks that meet the short detection definition are shown. Scale bar: 5 µm.

Pálfy, M., Schulze, G., Valen, E., & Vastenhouw, N. L. (2019). Chromatin accessibility established by Pou5f3, Sox19b and Nanog primes genes for activity during zebrafish genome activation. *BioRxiv*, 1–25. https://doi.org/10.1101/639302

Pérez-Rico, Y. A., Barillot, E., & Shkumatava, A. (2020). Demarcation of Topologically Associating Domains Is Uncoupled from Enriched CTCF Binding in Developing Zebrafish. *IScience*, *23*(5), 101046. https://doi.org/10.1016/j.isci.2020.101046
